# Evolutionary Coupling of Flagellar Motility and Type VI Secretion Systems Across Bacteria

**DOI:** 10.64898/2026.06.16.732483

**Authors:** Jamiema Sara Philip, Luke McNally, Matthew AB Baker

## Abstract

Bacteria use both motility and antagonism to compete in spatially structured environments, but whether these traits evolve together across broad bacterial diversity remains unclear. We developed a spatial kin-competition model predicting that motility should couple robustly with contact-dependent weapons by increasing encounter rates with competitors, whereas coupling with diffusible weapons should be weaker and context-dependent. To examine the evolutionary hypotheses motivated by the model, we performed a comprehensive analysis of 11,365 bacterial genomes across the bacterial Tree of Life (ToL). By utilising large-scale homology-based searches, we annotated flagellar, T6SS, and bacteriocin components and then applied phylogenetic comparative models to examine evolutionary associations. T6SS presence was strongly associated with flagellar motility: T6SS-positive lineages were predominantly flagellated, and BayesTraits supported a dependent model of FliC and T6SS evolution. In contrast, bacteriocins did not show detectable evolutionary coupling with flagellar motility. Transition-rate analyses further indicated that T6SS gain was strongly biased towards motile bacterial lineages, even though T6SS loss was common overall. These results indicate different macroevolutionary associations of flagellar motility with T6SS and bacteriocins, consistent with the qualitative hypothesis motivated by our model.

## Introduction

Natural environments host highly diverse microbial communities, in which bacteria compete intensively for limited resources such as nutrients, physical space, and oxygen (Hibbing et al. 2010). To survive in these settings, bacteria employ diverse ecological strategies, ranging from dispersal via motility to antagonistic interactions by diffusible toxins or contact-dependent weapons (Smith et al. 2023). Diffusible systems, such as bacteriocins, can act over relatively long distances, whereas contact-dependent systems such as the Type VI secretion system (T6SS) require direct physical contact to intoxicate competitor cells (Booth et al. 2023). The benefits of contact-dependent systems depend on the frequency of bacterial cell– cell encounters within microbial communities (Booth et al. 2023). Motility may therefore influence contact-dependent competition by increasing encounter rates between neighbouring cells and potential competitors (Tesseur et al. 2025).

Flagellar motility is a widespread and evolutionarily conserved form of bacterial motility (Miyata et al. 2020; Wadhwa and Berg 2022). Powered by the bacterial flagellar motor (BFM), it enables cells to navigate their environments and enhances dispersal, nutrient acquisition and colonisation (Nakamura and Minamino 2019; Philip et al. 2025).

Importantly, both flagella and T6SS are large multicomponent nanomachines encoded by conserved core structural genes that can be robustly identified across large genome datasets, making them especially suitable for broad-scale comparative evolutionary analyses. Both systems are energetically costly and require the coordinated expression of multiple tightly regulated genes (Alteri and Mobley 2016; Schavemaker and Lynch 2022). Since the macroevolutionary relationship between bacterial motility and bacterial warfare systems remains poorly understood, we quantified associations of a flagellar marker with T6SS-associated repertoires and BAGEL4 bacteriocin-associated hits.

Our recent large-scale comparative study revealed widespread gains and losses of flagellar systems across the bacterial tree of life, highlighting the dynamic evolutionary history of flagellar motility (Philip et al. 2025). Similarly, genomic surveys of the Type VI secretion system (T6SS) have uncovered substantial diversity in T6SS architecture, effector repertoires, and phylogenetic distribution across bacterial lineages (Unterweger et al. 2014; Lewis et al. 2019; Habich et al. 2025). Experimental disruption of T6SS components has also been associated with changes in motility and flagellar gene expression in several *Vibrio* species, although the mechanisms underlying these effects remain unclear (Wu et al. 2023; Lu et al. 2025). These observations remain largely restricted to mechanistic studies in individual organisms, and whether flagellar motility and T6SS exhibit coordinated evolutionary patterns across broad bacterial diversity and deep evolutionary timescales is yet to be established.

Whilst the T6SS is not only a contact weapon, its selective value as a bacterial weapon has a simple interference component: killing competitors reduces competitor survival and can relieve local competitive pressure (Chao and Levin 1981; Booth et al. 2023; Booth et al. 2024). For a contact-dependent system such as the T6SS, this benefit is encounter-limited, so motility should increase weapon value directly by increasing the number of susceptible competitors a producer contacts and can kill (Booth et al. 2023; Booth et al. 2024). Killing may also release resources or space. Because T6SS-mediated killing occurs at cell-to-cell distance, the producer is positioned to capture these benefits directly. Diffusible weapons such as bacteriocins create a different spatial problem: toxin-mediated killing can occur away from the producer, so benefits depend on whether the killed cell was competing locally, whether the producer can reach the freed patch, or whether nearby immune conspecifics capture it (Chao and Levin 1981; Kerr et al. 2002; Gardner et al. 2004; Driscoll and Pepper 2010). The latter generates inclusive-fitness benefits only when those conspecifics are close relatives (Hamilton 1964a; Hamilton 1964b; West et al. 2002; West et al. 2006). Thus, motility should robustly favour contact-dependent weapons through increased encounter and killing rates, whereas its effect on diffusible weapons should be weaker and more context-dependent because direct killing, resource capture, conspecific density, and relatedness can interact in opposing ways (Hamilton and May 1977; Taylor 1992; Nadell et al. 2016).

We formalised this logic in a spatial inclusive-fitness model and used the model to motivate a qualitative hypothesis that we examined across bacterial diversity (Fig. 1). The model predicted that flagellar motility should have a stronger effect on the benefits of contact-dependent weapons than on diffusible weapons, whose payoffs depend on additional ecological parameters (West et al. 2002; Gardner et al. 2004; Booth et al. 2023; Booth et al. 2024). We examined the evolutionary hypothesis using T6SS and bacteriocins as examples of contact-dependent and diffusible antagonism, respectively. To do this, we analysed 11,365 bacterial genomes, annotated flagellar, T6SS and bacteriocin components, and applied phylogenetic comparative models to examine correlated evolutionary dynamics. Together, this framework links bacterial motility, spatial competition and the evolution of antagonistic systems across the bacterial tree of life.

**Figure 1.**
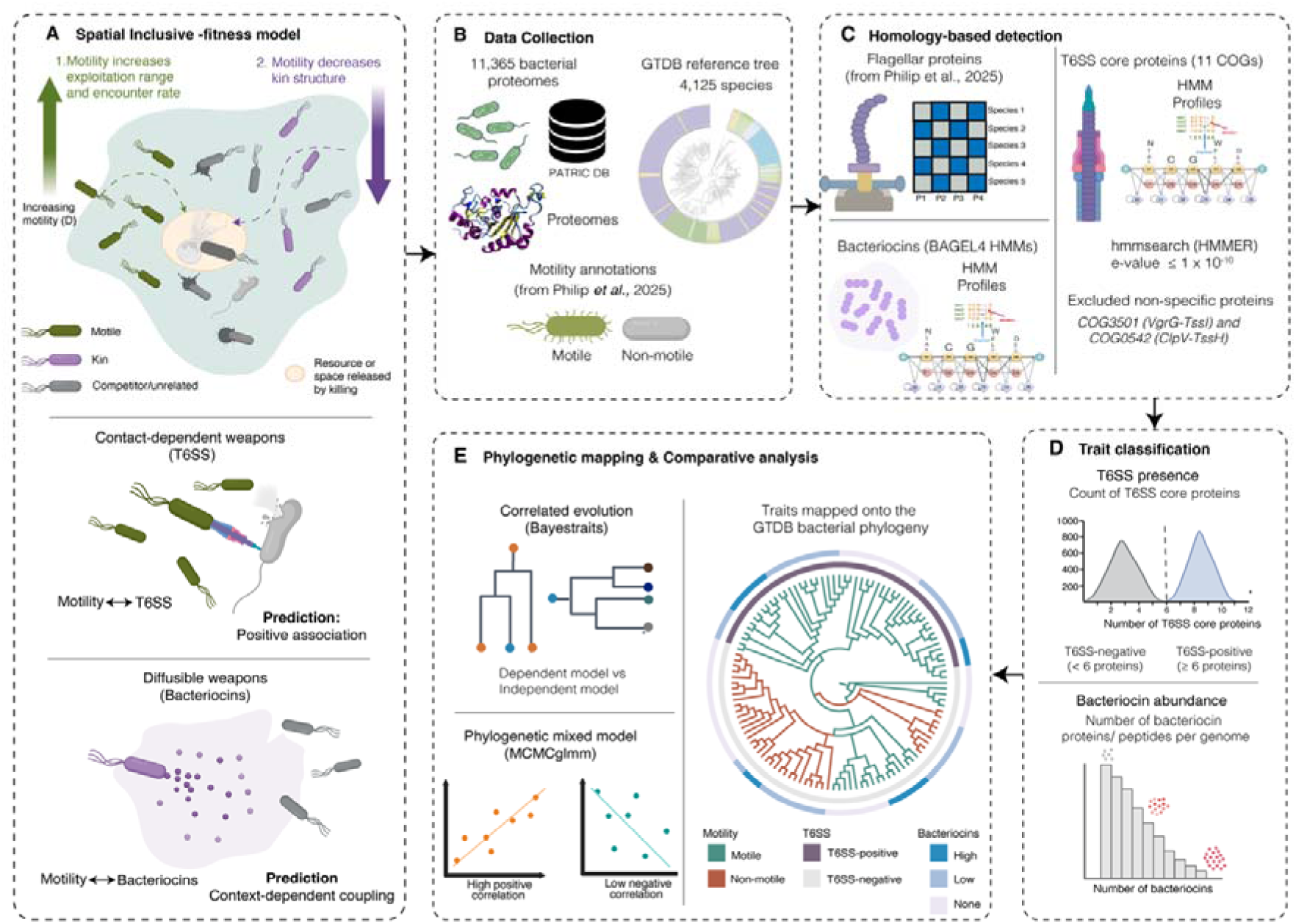
Theoretical framework and integrative workflow for analysing evolutionary associations between motility and bacterial competition systems. A spatial kin-competition theoretical framework predicting asymmetric coupling between motility and bacterial weapon classes was developed to guide comparative analyses. The model predicts a positive association between motility and contact-dependent systems such as the Type VI secretion system (T6SS), while predicting context-dependent associations with diffusible antagonistic systems such as bacteriocins through competing effects on exploitation range and local kin structure. A dataset of 11,365 bacterial proteomes and a reference bacterial phylogeny were then used to identify flagellar proteins, T6SS components and bacteriocins using homology-based searches (HMMER and BAGEL4). Motility annotations were obtained from a previously published dataset. T6SS presence and bacteriocin abundances were quantified per genome and encoded as binary or count-based traits. Traits were mapped onto the phylogeny and analysed using models of correlated evolution (BayesTraits) and phylogenetic mixed models (MCMCglmm). Selected icons in this figure were created using BioRender.com.

## Results

### Spatial kin-competition model predicts an asymmetric coupling between motility and the two weapon classes

The spatial kin-competition model generated different predictions for contact-dependent and diffusible weapons (Fig. 2). The model considers a focal weapon-producing cell surrounded by susceptible competitors and neighbouring conspecifics. Killing a competitor releases a resource or spatial opportunity that can be captured by the focal producer or nearby conspecifics, with the contribution of conspecifics to the producer’s fitness depending on their relatedness. Motility was represented as an increase in effective cell diffusion (D), affecting both access to released resources and the spatial structure of interactions. Increasing effective motility *D* expanded the producer’s exploitation reach *L*(*D*), allowing it to capture resources released by more distant kills, but also reduced the local relatedness scale ξ(*D*), representing the erosion of kin structure caused by mixing (Fig. 2A). For contact-dependent weapons, killing occurs at cell-to-cell distance and the producer is co-located with the released resource. As a result, the encounter-rate benefit of motility translated into a positive effect on contact-weapon fitness across the competitor densities examined (Fig. 2B). However, our model does not account for the duration of cell-to-cell contact, and at higher swimming speeds this benefit may diminish if cells separate before an attack can occur. Across parameter samples, the motility-induced change in contact-weapon fitness was tightly positive, motivating the hypothesis that motility may be strongly associated with T6SS-like weapons (Fig. 2F).

**Figure 2.**
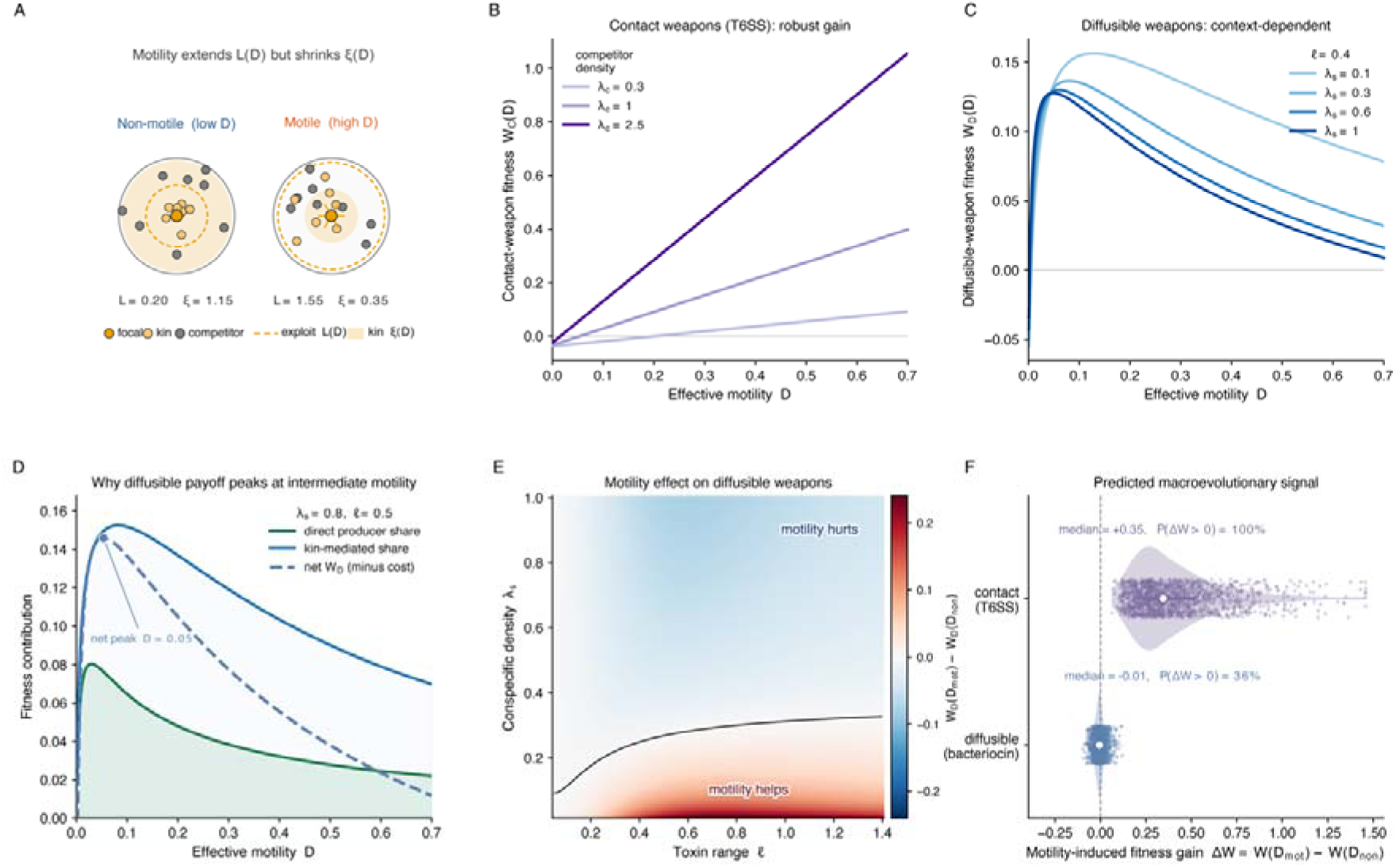
Spatial kin-competition model predicting asymmetric coupling between motility and bacterial weapon classes. **(A)** Motility increases the producer’s exploitation reach *L*(*D*), allowing it to access resources freed by killing competitors, but reduces the local relatedness scale ξ(*D*) by increasing spatial mixing. **(B)** For contact-dependent weapons, the model predicts that motility increases the rate of physical encounters with competitors, producing a consistently positive effect on weapon fitness across competitor densities. **(C)** For diffusible weapons, fitness depends on conspecific density because resources released by distant kills may be captured by either the producer or nearby immune conspecifics. **(D)** Decomposition of diffusible-weapon payoff into direct producer benefit and relatedness-weighted kin benefit. Both components can vary non-monotonically with motility. **(E)** Motility-induced change in diffusible-weapon fitness across toxin range and conspecific density. Positive values indicate regimes where motility favours diffusible weapons; negative values indicate regimes where motility disfavours them. **(F)** Parameter sampling shows that modelled motility effects are consistently positive for contact-dependent weapons but distributed around zero for diffusible weapons. Full equations, parameter values and sensitivity analyses are provided in Supplementary Methods S1, Supplementary Figs. S1–S4.

Diffusible weapons showed a more heterogeneous response. Because kills can occur away from the producer, their payoff depended on toxin range, conspecific density and the relatedness of conspecifics near the kill site. Decomposing the diffusible payoff showed that motility can increase the producer’s direct access to freed resources, but can also dilute indirect benefits by increasing conspecific competition and reducing local relatedness (Fig. 2C, D). Consequently, motility favoured diffusible weapons in some ecological regimes but disfavoured them in others, producing a sign change across toxin range and conspecific density (Fig. 2E). Across sampled parameter combinations, the motility-induced fitness change for diffusible weapons straddled zero (Fig. 2F). The model therefore predicted an asymmetric empirical signature: strong, signed coupling between motility and T6SS, but weak or heterogeneous coupling between motility and bacteriocins. We next examined these evolutionary hypotheses using comparative genomic and phylogenetic analyses.

### Global distribution of T6SS and bacteriocins across bacteria

We first quantified the genomic prevalence of T6SS and BAGEL-detected bacteriocins to determine whether these systems were sufficiently represented to test the model predictions. Across 11,365 bacterial genomes, we screened for core T6SS components and bacteriocin-associated proteins using homology-based searches. T6SS protein counts showed a bimodal distribution, with genomes either encoding few or no T6SS components or carrying a substantial complement of the system (Fig. 3A). Based on this distribution, species encoding six or more T6SS-associated proteins were classified as T6SS-positive, whereas species below this threshold were classified as T6SS-negative.

**Figure 3:**
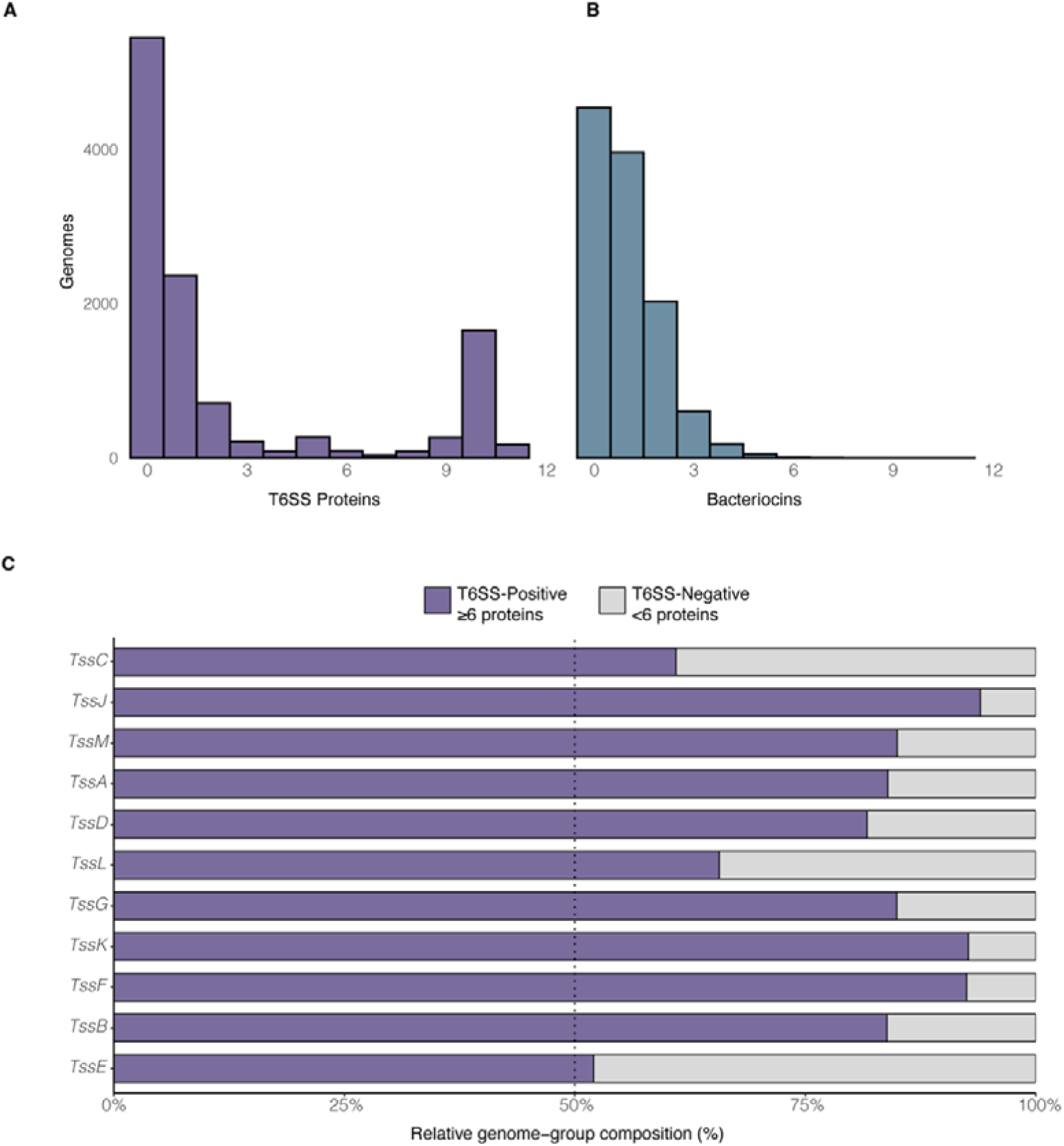
Distribution of Type VI secretion system (T6SS) proteins and bacteriocins across bacterial genomes. A) Histogram depicts the bimodal distribution of T6SS proteins. Genomes with fewer than six T6SS-associated proteins were classified as T6SS-absent, whereas genomes with six or more proteins were classified as T6SS-present, representing species likely to encode a putatively complete T6SS repertoire. The bimodal pattern highlights the discrete evolutionary distribution of T6SS across bacterial lineages. B) Histogram showing the distribution of bacteriocins across the same set of genomes. C) Stacked bar plot showing the distribution of individual T6SS core components (*TssA-TssM*) across the two groups. Genomes were grouped based on the T6SS protein content (<6 and ≥6 detected T6SS-associated proteins). The percentage of genomes which contained each T6SS protein was calculated within each group.

This threshold is consistent with previous research, which found that T6SS gene clusters with fewer than six core genes generally lack the essential components tssB and/or tssC (Morgado and Vicente 2022). These components are highly conserved and essential for T6SS function and classification, implying that smaller clusters are likely nonfunctional or degraded. We additionally examined the presence of these components among species classified as T6SS-positive. Of the 2,291 T6SS-positive species, 2,272 (99.2%) contained TssB, whereas 342 (14.9%) contained TssC.

Overall, 20.2% of species were classified as T6SS-present, while 79.9% were T6SS-absent. However, 52.1% of genomes have at least one T6SS protein. This distribution indicates that a putatively complete T6SS repertoire is restricted to a subset of bacterial lineages, with many species having only one or few T6SS proteins. The bimodal distribution highlights the discrete nature of T6SS acquisition and maintenance across the bacterial tree of life, with 18.3% of species having ≥9 core T6SS proteins. In contrast, bacteriocin-associated HMM hits showed a unimodal, right-skewed distribution, with most species having few or no hits (Fig. 3B).

To validate whether the six-protein threshold effectively captures genomes with a complete T6SS apparatus, we next examined the prevalence of individual core components (TssA– TssM) in genomes (Fig. 3C). Genomes classified as T6SS-positive (≥6 proteins) consistently exhibited a higher prevalence of every core component compared with T6SS-negative genomes. This confirms that the six-protein threshold enriches for genomes harbouring a more complete and potentially putatively complete T6SS repertoire. Among the small percentage of bacteriocin containing genomes (N≥4), we examined whether specific classes of bacteriocins dominate the diffusible antagonistic system repertoire. Class II bacteriocins (e.g., Bacteriocin IIc; 18.8%) were the most frequently detected, whereas Class I (e.g., Gallidermin; 1.2%) and Class III (e.g., Linocin_M18; 7.8%) bacteriocins were comparatively rare (Supplementary Fig. S5).

### Phylogenetic clustering of T6SS in motile bacterial lineages

We mapped T6SS presence alongside flagellar motility onto the Genome Taxonomy Database (GTDB) (Parks et al. 2022) bacterial phylogeny, using our previously published mapping framework (Philip et al. 2025). T6SS-positive species were concentrated in clades such as *Pseudomonadota*, *Desulfobacterota, Myxococcota, Acidobacteriota,* and *Planctomycetota (*Fig. 4). Notably, 87.9% of T6SS-positive species were classified as flagellar, whereas only 12.1% were classified as non-flagellar. Conversely, many non-flagellar phyla such as *Bacteroidota, Bacillota,* and *Actinobacteriota* lacked T6SS entirely. The phylogenetic distribution of these traits revealed a structured association between flagellar motility and T6SS across bacterial lineages. In contrast, bacteriocin showed no structured association with flagellar motility.

**Figure 4.**
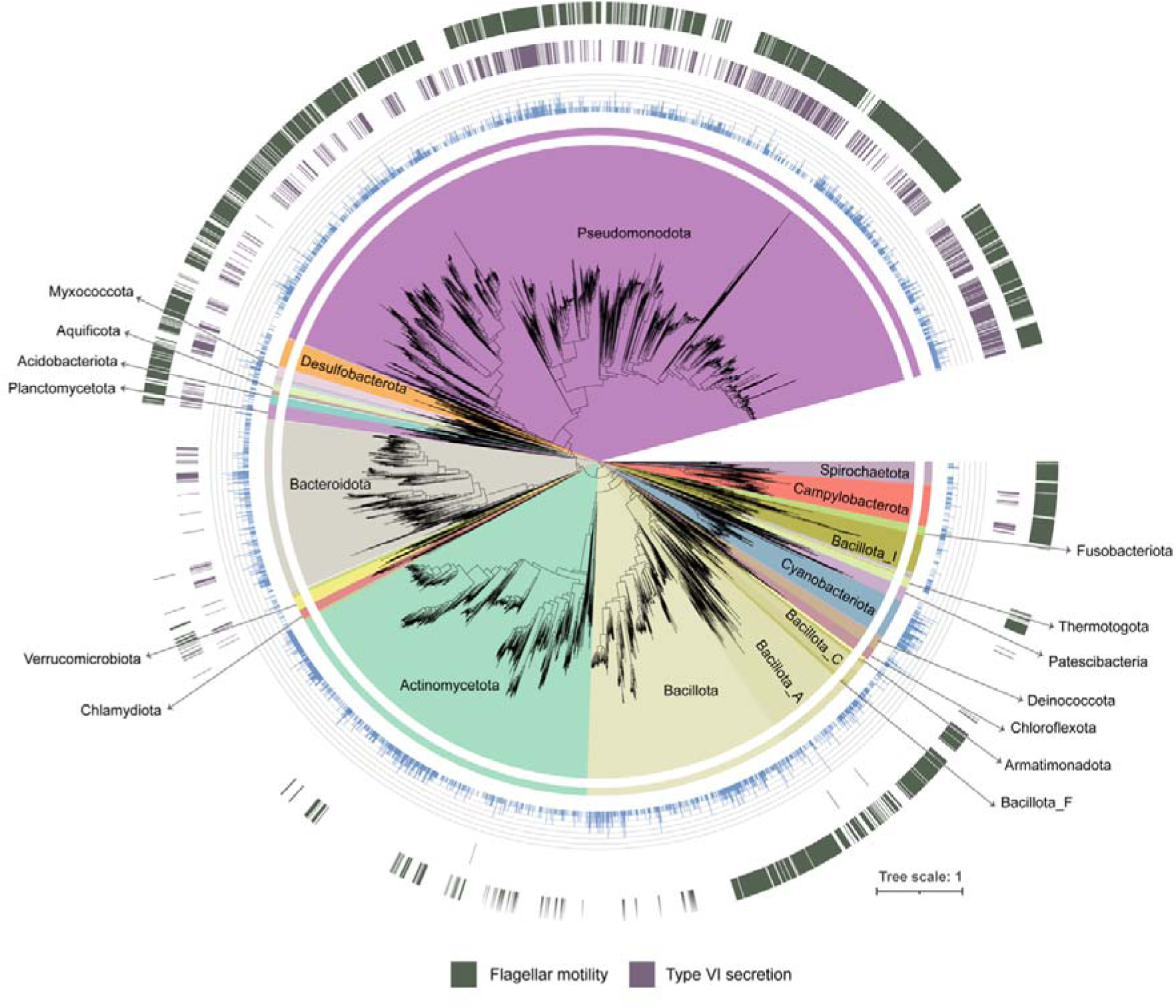
Phylogenetic distribution of T6SS and flagellar motility across the bacterial tree of life. The GTDB bacterial phylogeny (Philip *et al*., 2025) is shown at phylum-level resolution in a circular layout. The innermost ring indicates phylum-level classification, with colours distinguishing major bacterial groups. The inner bar plot indicates the bacteriocin count per species (scale 0-7, blue). The middle ring depicts T6SS presence, with T6SS-positive species in purple and T6SS-negative in white. The outer ring represents the presence of flagellar motility, with motile species in olive and non-motile in white.

### Co-occurrence of competition systems and flagellar proteins

To investigate the link between motility and inter-bacterial competition systems at the individual protein level, without accounting for phylogenetic non-independence, we examined first the simple co-occurrence of flagellar proteins with components of the Type VI secretion system (T6SS) and bacteriocin systems across bacterial genomes. Flagellar proteins showed a consistent positive pattern with T6SS components, with 97% of protein pairs exhibiting positive log odds ratios and a high average association strength (mean log OR = 2.61; median = 2.73) (Fig. 5A; Supplementary Table S1-S2). Positive log odds ratios were observed across multiple flagellar structural proteins and core T6SS components (e.g., FliG, FliM, FliN, FliC). In contrast, patterns between flagellar proteins and bacteriocins were weaker. Only 45.3% of protein pairs showed positive log odds ratios, while 54.7% had negative log odds ratios (mean log OR = -0.16; median = -0.12) (Fig. 5B; Supplementary Table S3-S4). Overall, T6SS components exhibited stronger and more consistent co-occurrence with flagellar proteins than bacteriocin systems across the datasets.

**Figure 5.**
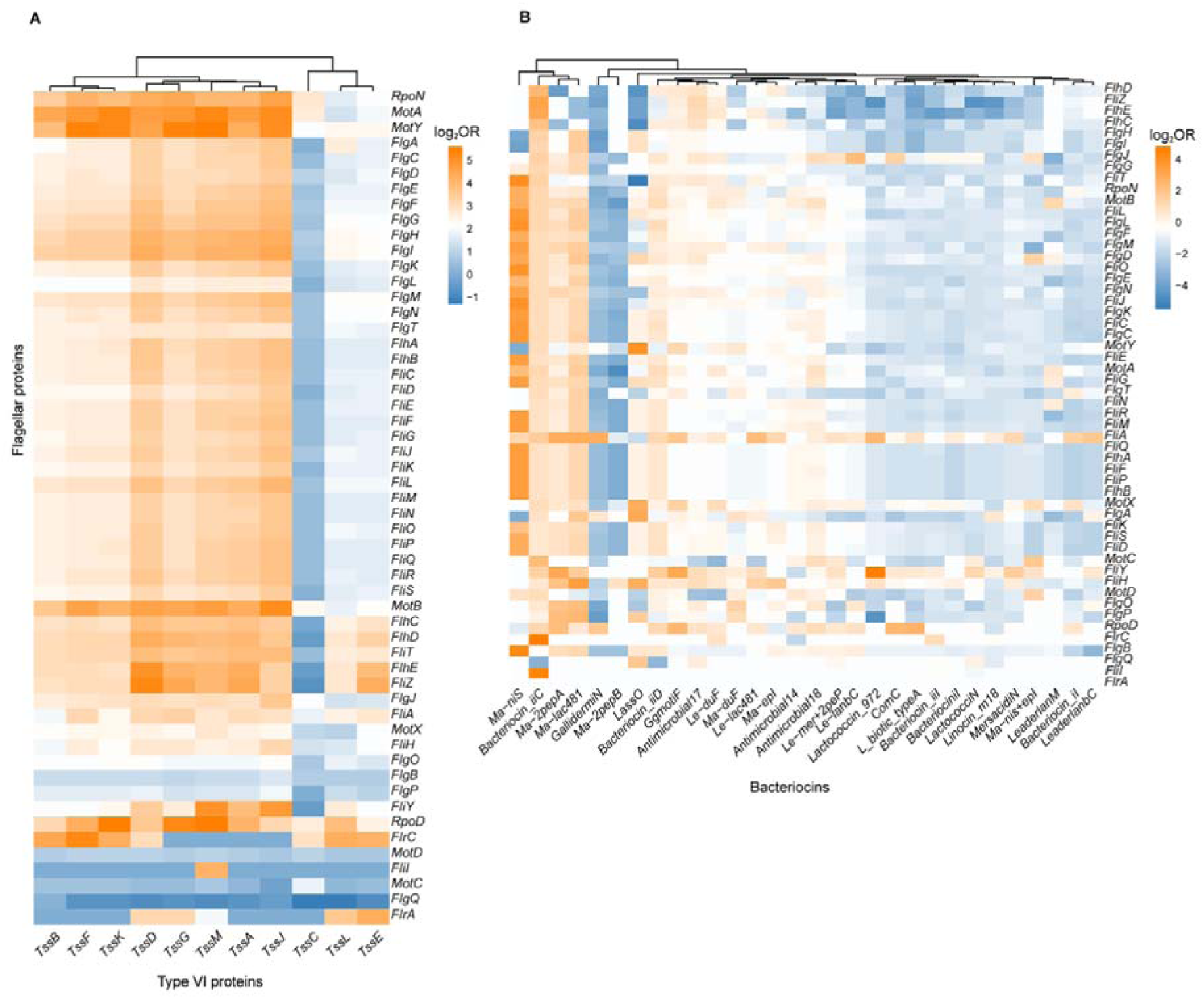
Protein-level co-occurrence between flagellar proteins and competition systems. Heatmap showing the co-occurrence patterns between flagellar proteins and (A) T6SS components, (B) Bacteriocins across bacterial genomes. Protein pairs were assessed using Fisher’s exact test based on protein presence–absence across bacterial species. Colours represent the log odds ratio, with orange indicating positive co-occurrence (proteins frequently found together in the same genome) and blue indicating negative co-occurrence.

### Evolutionary association between flagellar motility and T6SS

The most direct test of the model is the BayesTraits comparison of dependent and independent models of trait evolution, which asks whether motility and weapon presence have co-evolved across the bacterial phylogenetic tree. To test whether flagellar motility and competitive traits (T6SS or bacteriocin) evolved in a correlated manner, we applied phylogenetic comparative models (BayesTraits) comparing dependent and independent models of trait evolution. The analysis confirmed a strong evolutionary association between motility and T6SS with the dependent model yielding a log marginal likelihood of -2640.64 compared with -2668.15 for the independent model (Table 1). BayesTraits provided strong support for the dependent model of flagellar motility and T6SS evolution (Log BF = 55.03), indicating an evolutionary association between the two traits. Stochastic character mapping further reconstructed a median of 380 T6SS gains [95% CI: 285-511] in flagella-present lineages compared with 33 gains in flagella-absent lineages [95% CI: 10-67 (Supplementary Table S5). In contrast, the analysis provided no evidence of coupled evolution between motility and bacteriocin (log BF = -12.91).

**Table 1.** Evolutionary association between flagellar motility, T6SS and Bacteriocin: Log marginal likelihoods (Log ML) for the dependent and independent models of trait evolution calculated using BayesTraits.

| **Trait Comparison** | **Dependent Model (Log ML)** | **Independent Model (Log ML)** | **Log Bayes Factor** |
|---|---|---|---|
| FliC vs T6SS | -2640.64 | -2668.15 | 55.03 |
| FliC vs Bacteriocin | -1618.29 | -1608.15 | -12.91 |

### Evolutionary dynamics of flagellar motility and T6SS in bacteria

We estimated the transition rates between flagellar motility and T6SS states under the dependent model (Fig.6; Supplementary Fig. S6). Across bacterial lineages, transitions from T6SS positive to T6SS-negative states occurred at substantially higher rates than the gains of T6SS (q_21_ = 5.657; q_43_ = 6.391), indicating that T6SS loss is more common than T6SS gain during bacterial evolution. Posterior distributions of the transition-rate estimate further supported this asymmetry, with T6SS loss rates consistently exceeding gain rates across the posterior sample (Supplementary Fig. S6). However, the rate of T6SS gain was relatively higher in flagellar lineages (q_34_=1.316) than in non-flagellar lineages (q_12_ = 0.098). Together, these results indicate that although T6SS is frequently lost across bacterial evolution, its estimated gain rate is higher in flagellar than in non-flagellar lineages. This asymmetry in gain rates between flagellar and non-flagellar lineages (q_34/q_12 ≈ 13.4, Fig. 7) is consistent with the encounter-rate hypothesis proposed by the spatial model (Fig. 2B, F).

**Figure 6.**
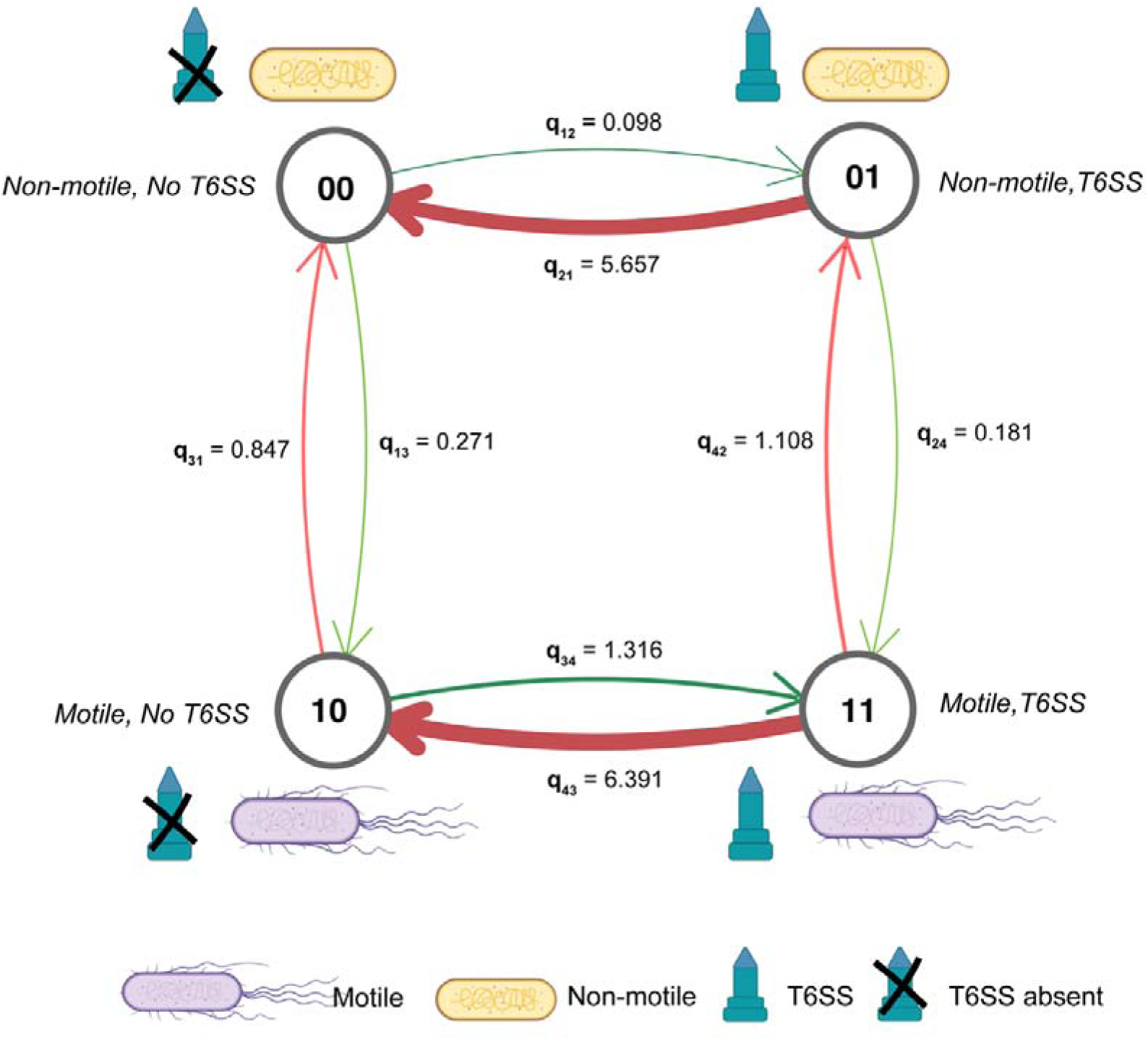
Evolutionary transition rates between motility and T6SS states inferred using BayesTraits. Transition rates between motility and T6SS states estimated under the dependent model of trait evolution in BayesTraits. The analysis evaluates transitions between four combined states representing motility (present or absent) and T6SS (present or absent). Selected icons in this figure were created using BioRender.com.

**Figure 7.**
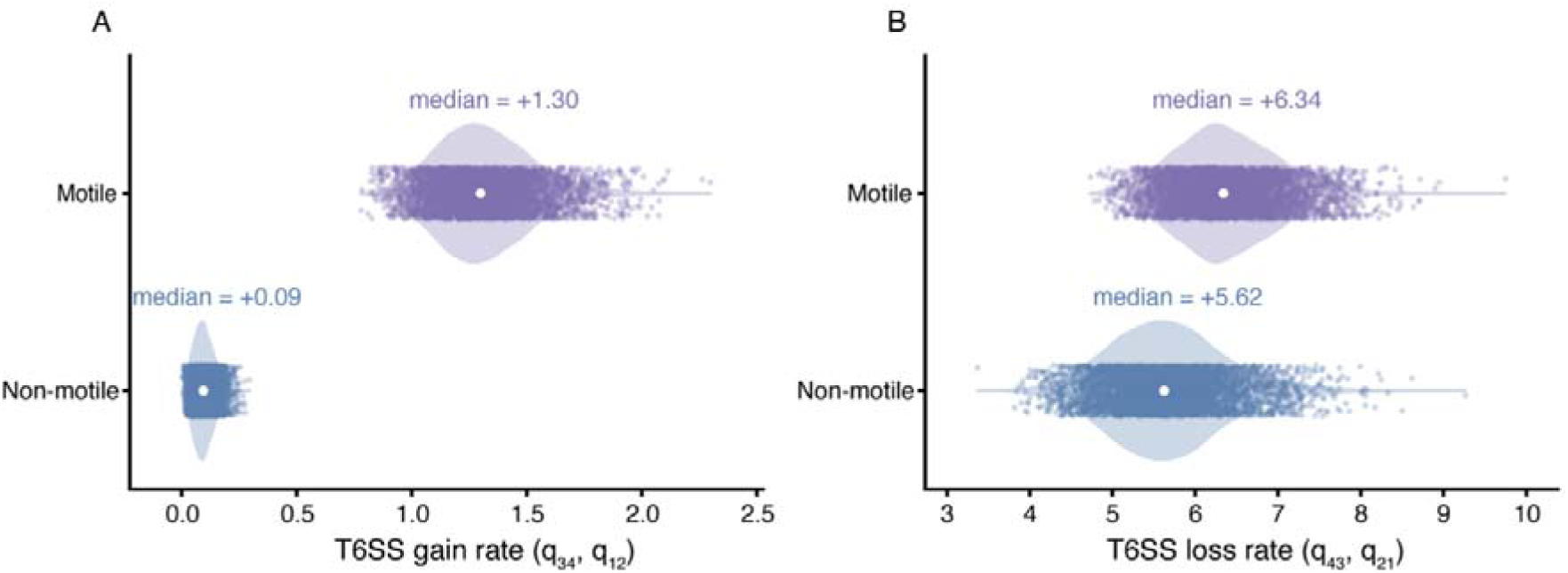
Posterior distributions of gain and loss of T6SS. Posterior distributions of gain and loss (B) of T6SS shown based on motility state as estimated in our BayesTraits analysis.

To explore the models in combination, we also fitted multivariate phylogenetic mixed models to quantify the phylogenetic relationships among motility, T6SS, and bacteriocin systems across bacterial diversity. Both flagellar motility and T6SS exhibited strong phylogenetic structure, with phylogenetic signal nearly 1 for FliC (median λ = 0.999, 95% credible interval (CI): 0.986–0.9999) and remaining high for T6SS (median λ = 0.866, 95% CI: 0.834–0.894) (Supplementary Fig. S8; Supplementary Table S6). These results indicate that closely related bacterial lineages maintain similar flagellar and T6SS states over time. In contrast, bacteriocins exhibit low phylogenetic signal (median λ = 0.039, 95% CI: 0.0002–0.175), indicating weaker phylogenetic structure in bacteriocin presence across the species analysed. Consistent with this pattern, posterior distributions of BayesTraits transition rates revealed substantially higher loss than gain rates for both T6SS and bacteriocins (Supplementary Figs. S7–S8), indicating frequent evolutionary turnover of antagonistic systems across bacterial genomes. However, despite this turnover, T6SS gains remained strongly associated with motile lineages, whereas bacteriocins exhibited substantially weaker long-term evolutionary stability and phylogenetic coupling.

Phylogenetic correlation analysis identified a positive evolutionary association between motility and T6SS (median correlation = 0.356, 95% CI: 0.211–0.508). This indicates that bacterial lineages retaining flagellar motility also tend to retain or encode more T6SS components over evolutionary time (Supplementary Table S7; Supplementary Fig. S8). In contrast, bacteriocins showed weak correlations with motility (median = 0.044, 95% CI: −0.594–0.577), while the correlation between T6SS and bacteriocins showed broad credible intervals overlapping zero (median = 0.527, 95% CI: −0.196–0.863). Consistent with these patterns, phylogenetic variance estimates were substantially higher for motility and T6SS traits than for bacteriocins, indicating stronger phylogenetic structure in motility and T6SS distributions across bacterial lineages (Supplementary Tables S8-9). These findings suggest a coupled evolutionary relationship between motility and T6SS, while bacteriocins exhibit more dynamic, lineage-independent evolutionary patterns.

Model diagnostics indicated good convergence for most T6SS and bacteriocin associated parameters, which showed high effective sample sizes and low autocorrelation across posterior samples (Supplementary Table S10-S11; Supplementary Fig. S9). However, several parameters involving motility showed poorer mixing, including lower effective sample sizes and elevated autocorrelation, particularly the FliC phylogenetic variance and the FliC–T6SS phylogenetic covariance estimates (Supplementary Table S11; Supplementary Fig. S9). Consequently, parameters involving FliC should be interpreted cautiously, and the MCMCglmm results are considered exploratory.

## Discussion

We investigated whether flagellar motility is differentially associated with T6SS and bacteriocins, as examples of contact-dependent and diffusible antagonistic systems. Our spatial kin-competition model predicts that motility should more consistently favour contact-dependent systems by increasing encounters with competitors, whereas its effect on diffusible antagonism is context-dependent. Across the 4,125 species included in the phylogenetic analysis, flagellar motility was strongly associated with T6SS presence, whereas no comparable association with bacteriocins were detected under our primary classification. For contact-dependent weapons, motility could increase the rate of physical encounters with competitors and therefore provide a direct route to increased weapon payoff. For diffusible weapons, motility can increase access to resources released by distant kills, but this effect can be offset by conspecific competition and the erosion of local kin structure.

### Evolutionary coupling between flagellar motility and T6SS

Phylogenetic analysis revealed a strong evolutionary association between flagellar motility and T6SS distribution across bacteria. Among T6SS positive species, 87.9% were flagellated, and T6SS was largely restricted to motile phyla such as *Pseudomonadota, Desulfobacterota,* and *Myxococcota*, while non motile phyla including *Bacteroidota, Bacillota,* and *Actinobacteriota* were T6SS negative (<6 T6SS core proteins) (Fig. 4). BayesTraits provided strong support for a dependent model of flagellar motility and T6SS evolution (log BF = 55.03; Table 1). Clade-restricted analyses showed that this association remained strongly supported within Pseudomonadota (log BF = 16.68), but not after excluding Pseudomonadota (log BF = -15.83), indicating that support is concentrated within this clade (Supplementary Table S12). This pattern was also reflected in the stochastic character mapping. Because stochastic mapping was performed separately for each clade-restricted dataset using its corresponding fitted model, the reconstructed transition counts are conditional on each analysis and are not subsets of the counts from the full dataset. Within Pseudomonadota, a median of 509 T6SS gains (95% CI: 380–684.5) were reconstructed in flagella-present lineages compared with 114 gains (95% CI: 65–197) in flagella-absent lineages (Supplementary Table S13). After excluding Pseudomonadota, similar numbers of T6SS gains were reconstructed in flagella-present lineages (mean = 18; 95% CI: 5–37) and flagella-absent lineages (median = 16; 95% CI: 5–30) (Supplementary Table S14).

At the individual protein level, flagellar and T6SS proteins showed a strong positive co-occurrence across genomes, even in the absence of phylogenetic information, with 8/11 T6SS essential proteins showing log_2_ odds ratios > 2 (Fig. 5A), suggesting a functional or regulatory link between these systems. Previous studies have shown that mutations in T6SS genes reduce flagellar gene expression and motility in species like *Citrobacter freundii* (Liu et al. 2015) *and Pseudomonas fluorescens* (Bouteiller et al. 2020). However, whether these effects reflect direct regulatory interactions between the two systems remains unclear. Shared regulatory architecture could also contribute to the observed association between flagellar motility and T6SS. For example, the alternative sigma factor σ (RpoN), which is involved in the regulation of flagellar genes in several bacterial lineages (Tsang and Hoover 2014), has also been shown to regulate T6SS gene clusters in diverse bacteria (Bernard et al. 2011). Such shared regulation could therefore contribute to the co-occurrence of flagellar motility and T6SS independently of or alongside selection arising from increased cell-cell encounters.

Bacteria use flagella and chemotaxis to disperse rapidly through heterogeneous environments, colonising favourable niches and encountering a diverse array of unrelated competitors along the way (Hibbing et al. 2010). Beyond the encounter-rate hypothesis illustrated in Fig. 2B, motility thus exposes producers to a phenotypically and phylogenetically diverse pool of competitors (Hamilton and May 1977), which may further amplify the selective advantage of a broadly targeting contact-dependent weapon such as T6SS (Codling et al. 2008; Nadell et al. 2016), although some bacteriocins can also exhibit broad target ranges (Cotter et al. 2013).

In structured communities such as biofilms, cells interact repeatedly with the same neighbours due to spatial structuring and clonal growth (Brockhurst 2007). Although these conditions provide frequent cell-cell contact that can favour T6SS activity, interactions may be largely restricted to local neighbours rather than the more diverse encounters potentially facilitated by motility. Thus, both the frequency and diversity of cell-cell encounters may influence the evolutionary benefits of T6SS.

Finally, our hypothesis does not require flagellar propulsion and T6SS firing to occur simultaneously. Motility may alter which organisms are encountered, whereas T6SS deployment could occur subsequently, once sufficiently close and sustained contact has been established. Swimming speed, encounter duration and the timing of T6SS deployment are not explicitly represented in the current model. At high movement rates, shorter contact times could reduce the probability of a successful T6SS attack.

### Energetic costs versus ecological utility

BayesTraits transition rate estimates (Fig. 6) revealed a strong asymmetry in T6SS evolution, with loss occurring more frequently than gain. Transitions from T6SS-positive to T6SS-negative states were higher (q = 5.66; q = 6.39) than the reverse (q = 0.098; q = 1.32), indicating that T6SS is evolutionary labile and was lost frequently across bacterial lineages. This pattern is consistent with broader evolutionary trends, in which loss of complex molecular systems is more common than their gain, including flagellar motility (Philip et al. 2025). However, the evolutionary dynamics of T6SS were associated with motility state. In non-motile lineages, gain of T6SS is rare (q = 0.098), whereas motile lineages show a higher probability of gaining T6SS (q = 1.32). Accordingly, T6SS loss exceeds gain by ∼58-fold in non-motile lineages, making acquisition highly rare, while motile lineages show a much weaker bias (∼5-fold). This suggests that although T6SS is generally prone to loss, its estimated gain is higher in motile bacterial lineages.

Flagellar systems impose a substantial energetic burden (∼10–17% of total cellular energy) (Schavemaker and Lynch 2022) and experimental deletion of ∼70 kb of flagellar genes in *Pseudomonas putida* eliminated swimming while improving growth efficiency (Martínez-García et al. 2014). In contrast, T6SS accounts for less than 1% of the ATP budget (Septer et al. 2023; Taillefer et al. 2023), yet our analysis indicates that T6SS is lost more frequently than flagellar motility (Fig. 6), suggesting that energetic cost alone does not explain its evolutionary dynamics. Instead, a major fitness benefit of T6SS arises from direct cell-cell competition, and its maintenance additionally requires coordinated structural, effector and immunity components (Andersson et al. 2017). T6SS function can vary with its effector repertoire, with some systems also involved in interactions with eukaryotic cells and nutrient acquisition (Alcoforado Diniz et al. 2015; Yang et al. 2021). As effector repertoires were not examined here, these functions may also contribute to the observed evolutionary association. In contrast, flagellar motility provides broader ecological functions, including chemotaxis, dispersal and colonisation, facilitating resource acquisition in heterogeneous environments (Mongold and Lenski 1996). It may be lost more frequently in host-associated environments (Sharp and Foster 2022), where host immune recognition of flagellin may impose additional selective pressure (Cullender et al. 2013).

### Alternative mechanisms of motility and antagonism

Although this study focuses on flagellar motility and T6SS, bacteria use other mechanisms of motility and antagonism that may also shape competitive interactions. For example, twitching and gliding motility can facilitate contact-dependent interactions (Booth et al. 2024; Lien et al. 2024), while CDI and T4SS provide alternative mechanisms of contact-dependent antagonism (Aoki et al. 2010; Czolkoss et al. 2021). Some bacterial lineages classified as non-flagellated particularly members of Bacteroidota, do exhibit gliding motility, highlighting a limitation of our specific focus on flagellar motility. These alternative systems for motility may contribute to broader evolutionary relationships between bacterial motility and competition.

### Bacteriocins and diffusible antagonism

The model prediction that the benefit of a diffusible weapon is context-dependent (Fig. 2E, F) is consistent with the lack of support for the dependent model (log BF = -12.91). There are notable differences in the molecular architecture of each system. T6SS is a multi-component macromolecular complex that requires a minimum set of core structural genes to form a functional apparatus (Fig. 3A). In contrast, bacteriocins are typically encoded by small, modular gene clusters consisting of only a few components (e.g., small precursor peptides, modification enzymes, immunity proteins and sometimes ABC transporters) (Smits et al. 2020). This simple genetic architecture of bacteriocins likely facilitates their horizontal gene transfer, increasing their mobility across genomes and complicating the reconstruction of their evolutionary history (Inglis et al. 2013).

The inferred relationship between flagellar motility and bacteriocin presence was sensitive to the threshold used to classify bacteriocin-positive species. At thresholds of ≥1, ≥2 and ≥3 bacteriocin-associated HMM hits, the dependent model was supported (log BF = 19.44, 36.85 and 31.67, respectively). In contrast, at the threshold of ≥4 HMM hits, which classified 3.0% of species as bacteriocin-positive, the independent model was supported (log BF = -12.91), as it was at the more stringent threshold of ≥5 HMM hits (log BF = -11.66) (Supplementary Table S15). Thus, the absence of an evolutionary association between flagellar motility and bacteriocin presence was not consistent across detection thresholds. Our results therefore indicate no detectable coupling under the primary ≥4 HMM-hit threshold, rather than a general absence of an association between these traits. Whilst we observed phylogenetic coupling between T6SS and motility, exploratory phylogenetic mixed models suggested differential relationships for T6SS and BAGEL-detected bacteriocins with flagellar motility.

### Methodological and data limitations

Several data-driven limitations should be considered. Homology-based trait classification provides a scalable and reproducible framework for identifying conserved multi-component systems across thousands of genomes. Our analysis of diffusible antagonism was restricted to bacteriocins identified using BAGEL4 HMMs and therefore does not capture the full diversity of diffusible antibacterial systems.

The binary classification of T6SS, supported by its bimodal distribution and previous studies, enables comparisons across species but oversimplifies the significant structural and regulatory diversity among T6SS subtypes. Notably, the diversity of T6SS effector repertoires makes them difficult to define and compare across bacterial lineages. Therefore, we excluded these components from our analysis to avoid confounding effects.

The association between flagellar motility and T6SS presence was concentrated within Pseudomonadota and was not supported when this clade was excluded. Although the association remained supported within Pseudomonadota, this analysis did not directly account for differences in cell-envelope architecture across bacterial lineages, which remains a limitation of this study.

Similarly, phylogenetic mixed models using MCMCglmm encountered difficulties with convergence when applied to the full dataset, reflecting the computational challenges associated with fitting complex models to extensive phylogenies. This analysis was therefore performed on a single random subset of 500 species, and limited mixing was observed for some FliC-related parameters. These results should therefore be considered exploratory rather than confirmatory. While the use of reduced datasets enhances model stability, it may compromise statistical power and increase uncertainty in parameter estimates. Limited ecological context, including habitat type and community structure, limits inference on the selective pressures shaping these traits. Future work integrating genomics with ecological and experimental data will be useful to test the causal hypotheses emerging from this evolutionary study.

### Bidirectionality in evolutionary relationship between T6SS and flagellar motility

Our framework is anchored in motility, as the last common bacterial ancestor is known to be motile (Coleman et al. 2021; Philip et al. 2025), suggesting that dispersal preceded the diversification of competitive strategies. However, the relationship may be bidirectional. T6SS-mediated killing can facilitate niche invasion and competitive displacement, generating ecological conditions that favour dispersal and colonisation. For example, *Salmonella enterica serovar Typhimurium* uses its T6SS to eliminate competing commensals, facilitating successful gut colonisation (Sana et al. 2016). Such dynamics raise the possibility that successful competition could, in turn, select for increased motility, reinforcing the association between these traits.

That said, our analyses offer conditional state-dependent estimates showing that transitions from non-motile to motile states occurred at lower estimated rates than transitions from motile to non-motile states, regardless of T6SS state (q_13_ = 0.271 vs q_31_ = 0.847 in T6SS-negative bacterial lineages; q_24_ = 0.181 vs q_42_ = 1.108 in T6SS-positive lineages). Non-flagellar forms of motility are largely restricted to a small number of clades, particularly within Bacteroidota.

Overall, our observations indicate that while flagellar motility represents a major and well-characterised mechanism, a broader range of motility strategies and the multiple selective pressures acting on them likely contributes to shaping competitive interactions. Integrating multiple forms of motility, and antagonistic systems, will be essential for a comprehensive understanding of how movement influences the evolution and deployment of bacterial weapons.

## Methods

### Spatial kin-competition model

Our comparative predictions were motivated by a spatial inclusive-fitness model, with full equations and sensitivity analyses provided in Supplementary Methods S1. In the model, flagellar motility is represented as an effective cell-diffusion coefficient D (Codling et al. 2008; Berg 2018). Increasing D affects two spatial scales: the exploitation length L(D), over which a producer can capture the resource or space released by killing a competitor, and the relatedness length ξ(D), over which neighbouring conspecifics remain close relatives (Hamilton 1964a; Hamilton 1964b; Taylor 1992; West et al. 2002). For a contact-dependent weapon, the fitness effect scales with the rate of physical encounters with susceptible competitors and with the producer’s direct access to the freed resource (Smoluchowski 1918; Berg and Purcell 1977; Booth et al. 2024). For a diffusible weapon, killing occurs over a distance distribution with length scale ℓ, so the fitness effect integrates over kill distance and combines the producer’s direct access with the relatedness-weighted benefit accruing through nearby immune conspecifics (Chao and Levin 1981; Driscoll and Pepper 2010; Allen et al. 2013; Nadell et al. 2016). The model is illustrative rather than fitted; parameter values were chosen to span plausible ecological regimes. The Python implementation used to generate the model figure is provided as Supplementary File S2.

### Data Collection and Homology Search

Motility presence–absence data were obtained from our previously published dataset (Philip et al. 2025), and analyses were performed using the same 11,365 bacterial proteomes and the 4125 species GTDB (Parks et al. 2022) reference bacterial tree as the phylogenetic framework.

T6SS core proteins and bacteriocins were identified across the 11,365 bacterial proteomes using homology-based searches (Fig. 1). Proteomes were screened with hmmsearch (Finn et al. 2011) (e-value ≤ 1 × 10 ¹ ) against HMM profiles for 11 T6SS core COGs (Morgado and Vicente 2022), excluding COG3501 (*VgrG-TssI*) and COG0542 (*ClpV-TssH*), which are not T6SS-specific, and against bacteriocin HMM profiles from BAGEL4 (van Heel et al. 2018). Species encoding six or more of the screened T6SS-associated proteins were classified as T6SS-positive, whereas species below this threshold were classified as T6SS-negative.

For bacteriocins, the number of hits against BAGEL4 bacteriocin HMM profiles was counted for each proteome. These profiles represent bacteriocin-related sequence features, including bacteriocin families, leader peptides and other bacteriocin-associated motifs. Species with ≥4 detected HMM hits were classified as bacteriocin-positive, whereas species below this threshold were classified as bacteriocin-negative. To assess the effect of this classification threshold, the analysis was repeated using alternative cut-offs (≥1, ≥2, ≥3, ≥4 and ≥5) for bacteriocin presence.

### Phylogenetic trait analysis and correlated evolution

To enable comparative analysis across 11,365 bacterial genomes, we defined motility and bacterial weapon strategies as binary traits based on protein presence-absence.

The flagellar motility (FliC) presence–absence trait was obtained from our previously published dataset (Philip et al. 2025), in which genomes were classified as motile or non-motile based on the number of flagellar protein-encoding genes they contain. FliC, an established marker of flagellar motility, was used as the gene marker for the motility trait in the subsequent analyses.

The presence of a Type VI secretion system (T6SS) was inferred using a homology-based approach. Predicted proteomes were screened for canonical T6SS components, and the number of detected T6SS-associated proteins was quantified for each species. Across all genomes, the distribution of T6SS protein counts was bimodal, with one peak corresponding to species lacking T6SS components and the other species encoding a substantial complement of the system. Based on this bimodal distribution, species encoding six or more T6SS-associated proteins were classified as T6SS-positive, whereas species below this threshold were classified as T6SS-negative. This cutoff was chosen to distinguish genomes likely to encode a putatively complete T6SS repertoire from those containing partial or degraded homologues and has been used in previous studies to identify putatively complete T6SS gene clusters (Morgado and Vicente 2022).

The presence or absence of diffusible antagonistic systems was assessed using a similar homology-based screening approach. For each genome, the number of bacteriocin-associated HMM hits was recorded. Species with N ≥ 4 bacteriocin-associated HMM hits were classified as bacteriocin-present, while species with N < 4 were classified as bacteriocin-absent.

Finally, all traits (motility, T6SS, bacteriocins) were treated as discrete binary characters, with state 0 indicating absence and state 1 indicating presence for downstream phylogenetic comparative analysis.

To test for evolutionary dependence between flagellar motility and secretory traits (T6SS or bacteriocins), we used the Discrete model implemented in BayesTraits v4.0 (Pagel and Meade 2006), fitting both dependent and independent models of trait evolution and estimating their marginal likelihoods to assess model support. To assess the effect of broad taxonomic structure on the association between flagellar motility and T6SS presence, we repeated the analysis within Pseudomonadota and after excluding Pseudomonadota.

To estimate the number and type of evolutionary transitions between flagellar motility and type VI secretion system (T6SS) states, we performed stochastic character mapping conditional on the fitted BayesTraits dependent model. FliC presence/absence and T6SS presence/absence were represented as a single four-state character (00, 01, 10 and 11), corresponding to the four possible combinations of the two binary traits. For each posterior sample from the BayesTraits analysis, we constructed the corresponding continuous-time Markov transition matrix from the eight estimated transition rates and generated a stochastic character history on the phylogeny conditional on the observed tip states. We then counted the realised number of each of the eight permitted transition types across the tree. In particular, transitions from 00→01 represent gains of T6SS in lineages lacking flagella, whereas transitions from 10→11 represent gains of T6SS in lineages possessing flagella. Repeating this procedure across posterior samples allowed uncertainty in both model parameters and ancestral evolutionary histories to be propagated into posterior distributions for the number of each transition type. We additionally recorded the total branch length spent in each joint state to describe the evolutionary opportunity for each class of transition. To examine whether the reconstructed transition patterns were concentrated within Pseudomonadota, we repeated the stochastic character mapping within Pseudomonadota and after excluding Pseudomonadota from the phylogeny, using the corresponding clade-specific BayesTraits posterior transition rates. Analyses were implemented in R using ape and phytools.

### Co-occurrence of competition systems and flagellar proteins

To analyse the interplay between bacterial motility and competition strategies, the co-occurrence of flagellar proteins with Type VI secretion system (T6SS) and bacteriocins across various bacterial species was examined. A presence-absence dataset was created for all proteins from three systems across the species. Each flagellar and competition-related protein pair was assessed using a 2 × 2 contingency table to examine their co-occurrence. Fisher’s exact test was applied to each protein pair, and odds ratios were calculated to estimate the strength of co-occurrence. These odds ratios were subsequently log -transformed to allow for comparative analysis across different protein pairs. The final log odds ratios were visualised as heatmaps, highlighting co-occurrence patterns between flagellar proteins and components of the T6SS and bacteriocin systems. These analyses were treated as descriptive because they do not account for phylogenetic non-independence among species.

### Phylogenetic mixed model analysis using MCMCglmm

As an exploratory complementary analysis of the evolutionary relationships among motility, type VI secretion system (T6SS), and bacteriocins, we fitted a multivariate phylogenetic mixed model using a Bayesian framework implemented in MCMCglmm (Hadfield 2010). The dataset comprised species-level measures of FliC presence (binary), T6SS protein counts, and bacteriocin counts. Motility was modelled as a binary trait using a threshold model. T6SS was modelled as a binomial-type response, representing the number of detected core proteins relative to the total possible proteins (n = 11), while bacteriocin abundance was modelled as count data using a Poisson distribution.

To ensure computational tractability, a single random subset of 500 species was selected using a fixed seed (set.seed(123)) for reproducibility. The phylogenetic tree was pruned to include only the sampled taxa, and species names were matched between the dataset and the phylogeny. Phylogenetic relationships were incorporated via an inverse relatedness matrix derived from the pruned tree using the inverseA function in MCMCglmm.

A trivariate model was fitted with an unstructured variance–covariance matrix for both phylogenetic and residual effects, allowing estimation of covariance among traits. Weakly informative inverse-Wishart priors were specified for both variance components, with the residual variance for the threshold trait fixed for identifiability. Models were run for 500,000 iterations with a burn-in of 100,000 and a thinning interval of 500. Convergence was assessed through visual inspection of trace plots and by calculating effective sample sizes and autocorrelation diagnostics using the coda.

Phylogenetic correlations between traits were calculated from the posterior distributions of the variance–covariance matrix and reported as medians with 95% credible intervals. Trait-specific phylogenetic signal (λ) was estimated as the proportion of total variance explained by phylogeny.

## Data availability

All the data used are presented in the Supplementary Information.

## Code availability

The code used is available at https://github.com/BakerLabBABS/classifier/

## Author information

### Contributions

Jamiema Sara Philip (Investigation, Visualization, Formal Analysis, Writing—original draft, Writing—review & editing), Luke McNally (Investigation, Formal Analysis, Visualization, Writing—review & editing), Matthew AB Baker (Investigation, Formal Analysis, Conceptualization, Supervision, Project administration, Visualization, Writing—original draft, Writing—review & editing).

