## Supplementary Figs S1-S9, Supp Tables S1-S15 for "Evolutionary Coupling of Flagellar Motility and Type VI Secretion Systems Across Bacteria"

**Supplementary information for "Evolutionary Coupling of Flagellar Motility and Type VI Secretion Systems Across Bacteria"**

Jamiema Sara Philip, Luke McNally, and Matthew AB Baker

### SUPPLEMENTARY FIGURES

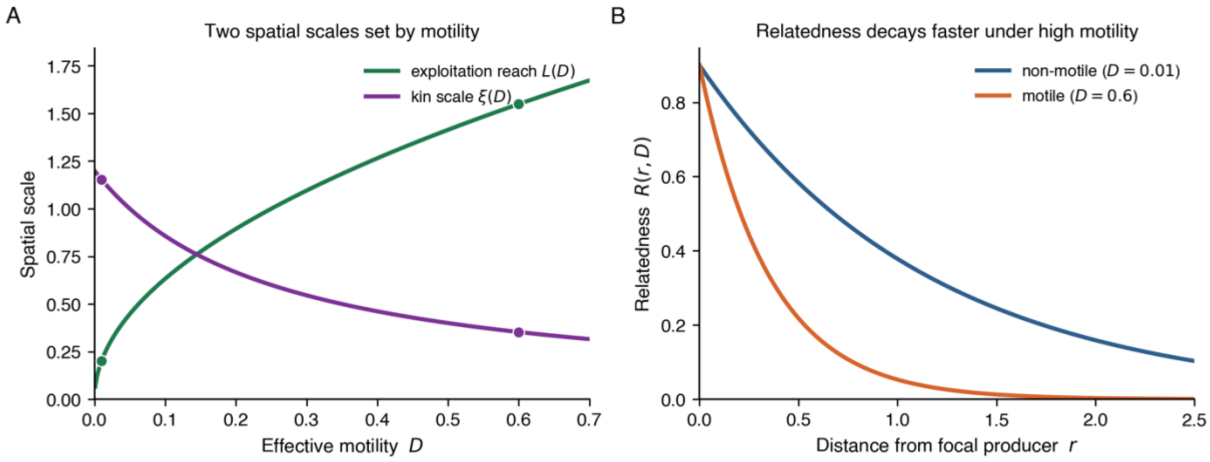


**Supplementary Figure S1.** Two competing spatial effects of motility, shown as functions of the effective diffusion coefficient D. (A) Exploitation reach L(D) (green) rises monotonically with D while the kin scale ξ(D) (purple) declines; filled markers indicate the non-motile (D = 0.01) and motile (D = 0.6) operating points used elsewhere. (B) Relatedness R(r, D) as a function of distance from the focal producer for the two operating points: high motility produces a steeper decay because ξ(D) is smaller.

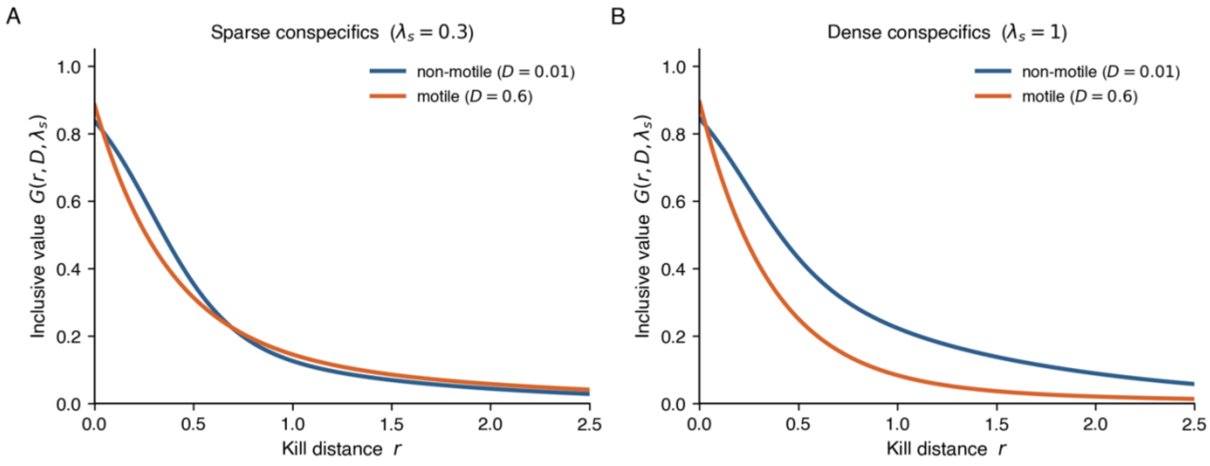


**Supplementary Figure S2.** Inclusive-fitness value G(r, D, λ_s) of a kill as a function of kill distance r, for non-motile (blue) and motile (orange) producers. (A) Sparse conspecifics (λ_s = 0.3): the two curves are similar because neither direct-access dilution nor kin contribution is strong. (B) Dense conspecifics (λ_s = 1.0): the non-motile producer extracts substantially more inclusive value at intermediate kill distances because the dense neighbourhood is filled with closer relatives (larger ξ(D)). Motility flattens both effects.

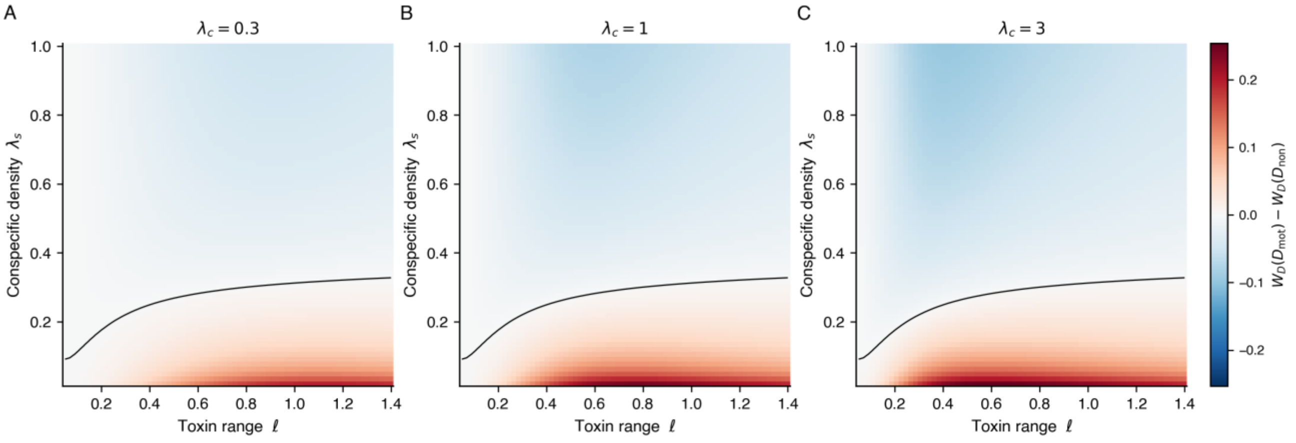


**Supplementary Figure S3.** Robustness of the diffusible-weapon sign-flip surface across competitor densities. Each panel shows W_D(D_motile) − W_D(D_non-motile) over toxin range ℓ and conspecific density λ_s, computed under (A) λ_c = 0.3, (B) λ_c = 1.0, and (C) λ_c = 3.0. The shape of the zero-contour (black) is essentially independent of competitor density; what changes is the magnitude of the motility effect. The prediction that motility favours diffusible weapons in the sparse, short-to-intermediate-ℓ regime and disfavours them elsewhere is therefore not an artefact of any particular λ_c.

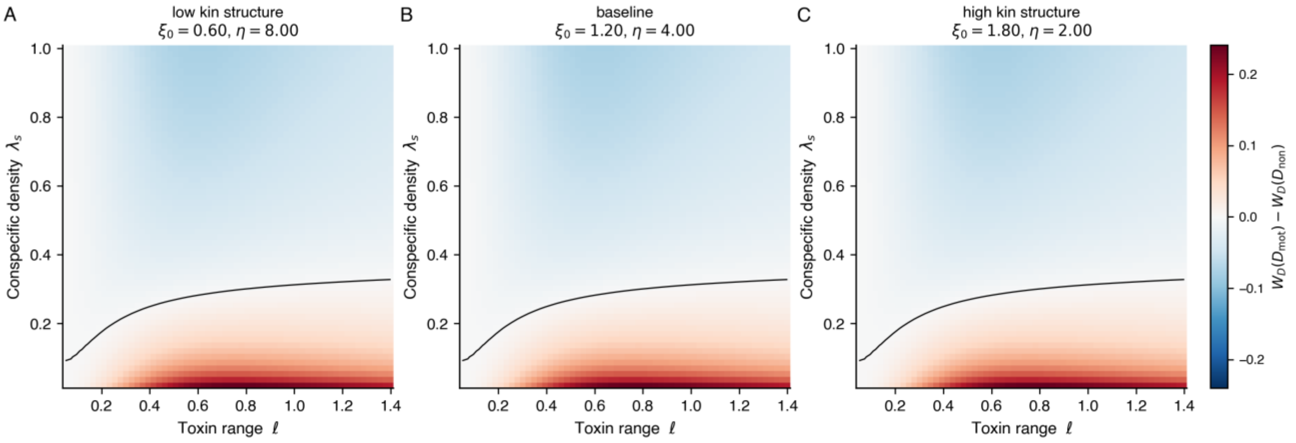


**Supplementary Figure S4.** Sensitivity of the diffusible-weapon motility effect to the relatedness parameters ξ_0 (baseline kin scale) and η (motility-driven kin erosion). (A) Low kin structure (ξ_0 halved, η doubled), (B) baseline (ξ_0 = 1.20, η = 4.00, as in Fig. 1), (C) high kin structure (ξ_0 increased, η halved). The qualitative position of the sign-flip surface is preserved across all three configurations, but the magnitude of the motility effect grows with the strength of kin structure. This confirms that the bacteriocin context-dependence predicted by the model depends on the existence of motility-mediated kin erosion, not on the precise functional form chosen here.

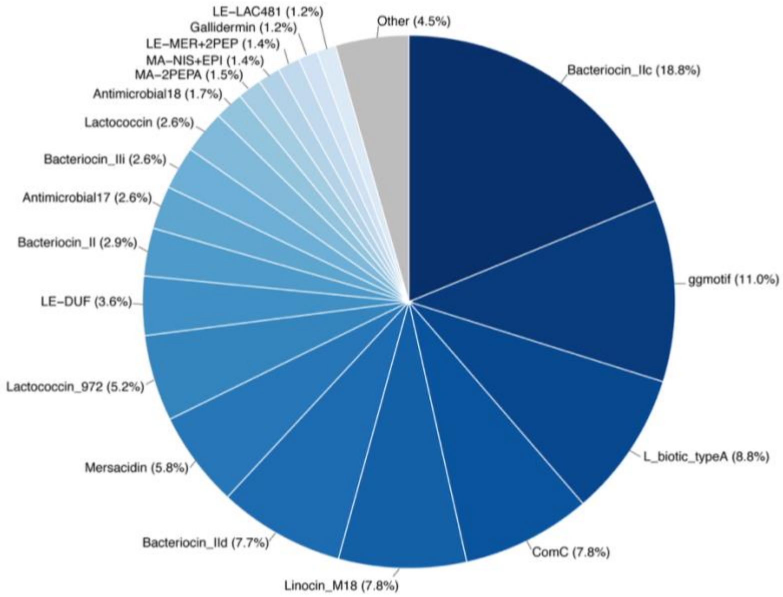


**Supplementary Figure S5. Distribution of bacteriocin classes across bacterial genomes.** Pie chart showing the relative abundance of different bacteriocin types identified across genomes.

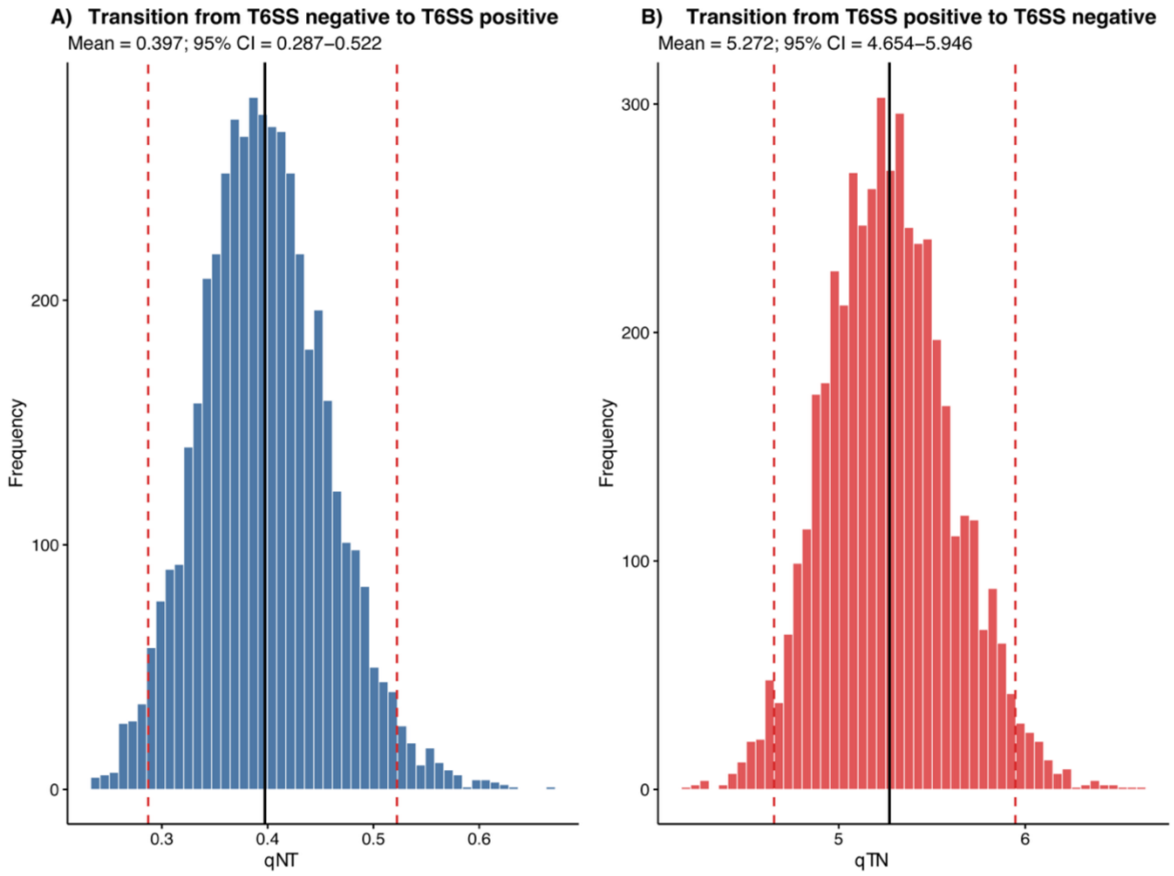

**Supplementary Fig S6. Bayestraits transition rates of T6SS.** Histograms indicate the posterior distribution of transition rate estimates between T6SS absent (qNT) and T6SS present (qTN). The black line indicates the mean, and the red dotted line indicates the 95% CI.

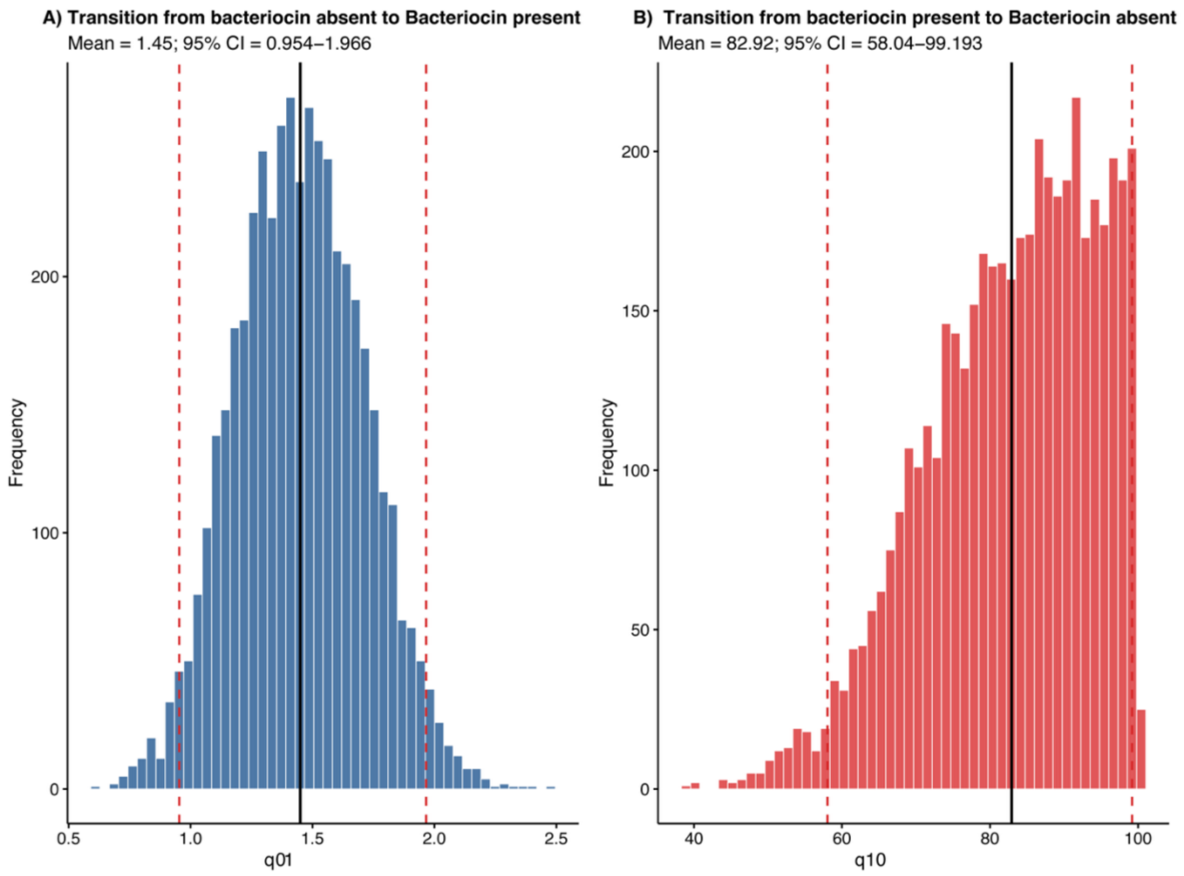


**Supplementary Fig S7. Bayestraits transition rates of Bacteriocin.** Histograms indicate the posterior distribution of transition rate estimates between Bacteriocin absent (q01) and Bacteriocin present (q10). The black line indicates the mean, and the red dotted line indicates the 95% CI.

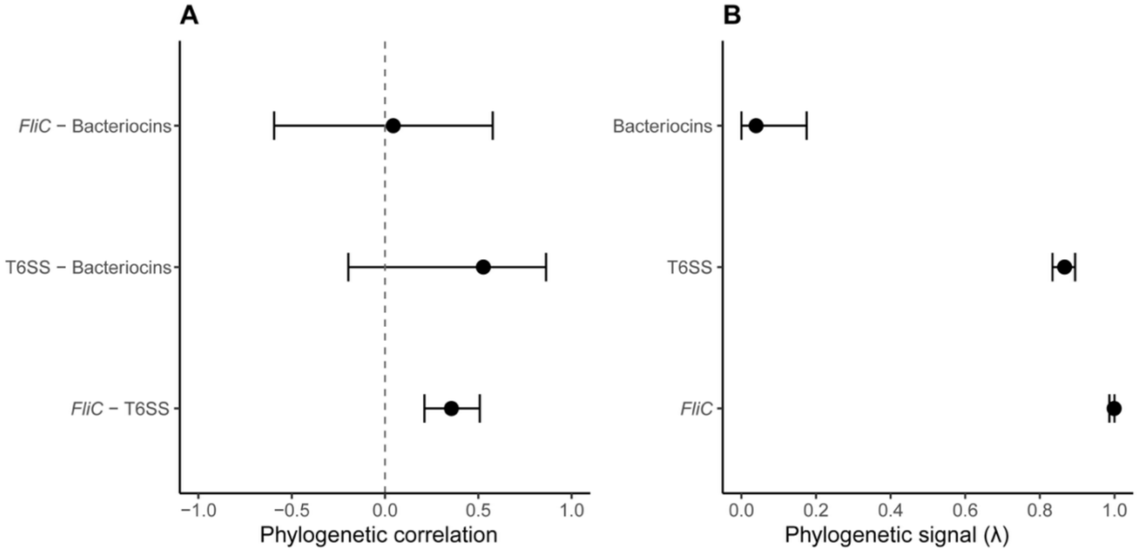


**Supplementary Figure S8. Phylogenetic correlations and signal of motility, T6SS and bacteriocins.** (A) Posterior estimates of phylogenetic correlations among flagellar motility (*FliC*), T6SS and bacteriocin abundance, inferred using a trivariate phylogenetic mixed model. Points represent posterior medians and horizontal bars indicate 95% credible intervals. The dashed vertical line denotes zero correlation. (B) Phylogenetic signal for each trait estimated as the proportion of total variance explained by phylogeny. Points represent posterior medians and horizontal bars indicate 95% credible intervals.

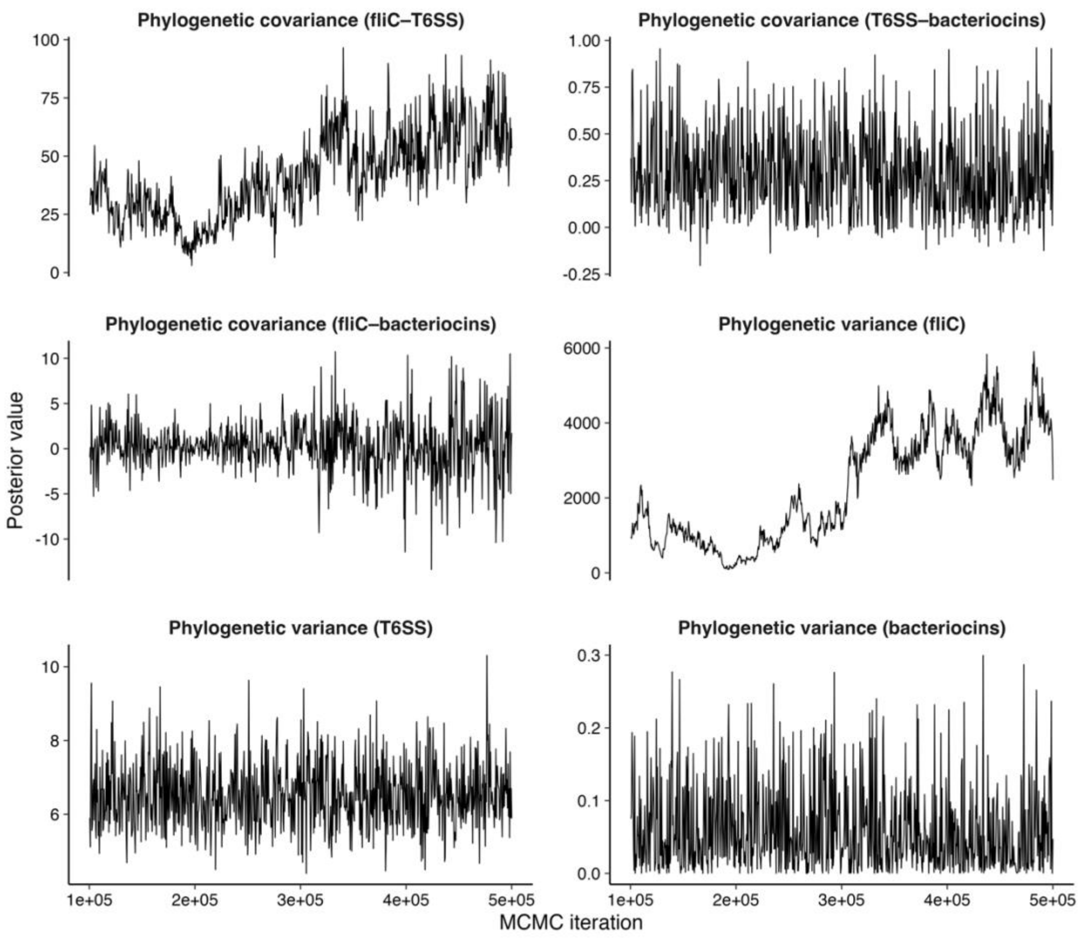


**Supplementary Figure S9: MCMC convergence diagnostics**. Trace plots showing posterior samples for phylogenetic variance and covariance components from the trivariate phylogenetic mixed model. Values are plotted against MCMC iteration number following burn-in and thinning. Panels correspond to variance components for each trait and covariance between trait pairs.

### Supplementary Tables

**Supplementary Table S1**. Summary statistics for pairwise co-occurrence analyses between flagellar and Type VI secretion system (T6SS) proteins across bacterial genomes.

**Supplementary Table S2. Results of pairwise co-occurrence analyses between flagellar and Type VI secretion system (T6SS) proteins.** Associations were quantified using odds ratios calculated from protein presence–absence patterns across bacterial genomes. The table includes odds ratios, p-values, FDR-adjusted p-values, $\log_2$-transformed odds ratios, and significance status following FDR correction (FDR < 0.05).

**Supplementary Table S3.** Summary statistics for pairwise co-occurrence analyses between flagellar and bacteriocins across bacterial genomes.

**Supplementary Table S4. Results of pairwise co-occurrence analyses between flagellar and bacteriocins**. Associations were quantified using odds ratios calculated from protein presence–absence patterns across bacterial genomes. The table includes odds ratios, p-values, FDR-adjusted p-values, $\log_2$-transformed odds ratios, and significance status following FDR correction (FDR < 0.05).

| Transition | State of other trait | Mean | SD | Median | 95% CI |
|---|---|---|---|---|---|
| T6SS gain | Flagella absent | 34.6 | 14.8 | 33 | 10–67 |
| T6SS gain | Flagella present | 384.7 | 57.0 | 380 | 285–511 |
| T6SS loss | Flagella absent | 180.5 | 22.2 | 181 | 136–225 |
| T6SS loss | Flagella present | 574.2 | 52.0 | 571 | 479–686 |
| Flagella gain | T6SS absent | 97.1 | 9.0 | 97 | 80.5–115 |
| Flagella gain | T6SS present | 4.7 | 3.3 | 4 | 0–13 |
| Flagella loss | T6SS absent | 247.3 | 15.8 | 247 | 216–279 |
| Flagella loss | T6SS present | 98.7 | 12.2 | 98 | 76–124 |

**Supplementary Table S5. Stochastic character mapping of transitions between flagellar motility and T6SS presence across the bacterial phylogeny.** The mean, standard deviation (SD), median and 95% interval of the number of reconstructed gains and losses are shown across stochastic character maps. T6SS gains and losses are reported according to whether flagella were present or absent, and flagellar gains and losses according to whether T6SS was present or absent.

| Trait | Median λ | Lower 95% CI | Upper 95% CI |
|---|---|---|---|
| *FliC* | 0.999 | 0.986 | 0.9999 |
| T6SS | 0.866 | 0.834 | 0.894 |
| Bacteriocins | 0.039 | 0.0002 | 0.175 |

**Supplementary Table S6. Trait-specific phylogenetic signal (λ):** Trait-specific phylogenetic signal (λ) estimated as the proportion of total variance explained by phylogeny.

| Trait comparison | Median correlation | Lower 95% CI | Upper 95% CI |
|---|---|---|---|
| *FliC* – T6SS | 0.356 | 0.211 | 0.508 |
| *FliC* – bacteriocins | 0.044 | -0.594 | 0.577 |
| T6SS – bacteriocins | 0.527 | -0.196 | 0.863 |

**Supplementary Table S7. Phylogenetic correlations among motility, T6SS, and bacteriocins:** Posterior phylogenetic correlations calculated from the phylogenetic variance–covariance matrix. Positive values indicate that traits tend to co-evolve across bacterial lineages.

| Phylogenetic variance/covariance | Posterior mean | Lower 95% CI | Upper 95% CI | Effective sample size |
|---|---|---|---|---|
| *FliC* | 2307.584 | 114.079 | 4658.875 | 2.78 |
| T6SS-*FliC* | 41.238 | 11.448 | 76.200 | 4.39 |
| Bacteriocins-*FliC* | 0.312 | -5.583 | 6.105 | 586.69 |
| *FliC*-T6SS | 41.238 | 11.448 | 76.200 | 4.39 |
| T6SS | 6.542 | 4.881 | 8.260 | 800.00 |
| Bacteriocins-T6SS | 0.291 | -0.055 | 0.751 | 800.00 |
| *FliC*-Bacteriocins | 0.312 | -5.583 | 6.105 | 586.69 |
| T6SS-Bacteriocins | 0.291 | -0.055 | 0.751 | 800.00 |
| Bacteriocins | 0.061 | 0.000 | 0.183 | 800.00 |

**Supplementary Table S8. Phylogenetic variance–covariance matrix (G structure) from the multivariate phylogenetic mixed model:** Posterior estimates of phylogenetic variances and covariances obtained from the multivariate phylogenetic mixed model fitted in MCMCglmm. Diagonal elements represent phylogenetic variances for each trait, whereas off-diagonal elements represent phylogenetic covariances between traits.

| Residual variance/covariance | Posterior mean | Lower 95% CI | Upper 95% CI | Effective sample size |
|---|---|---|---|---|
| *FliC* | 2.706 | 0.342 | 8.168 | 658.76 |
| T6SS-*FliC* | 0.074 | -1.188 | 1.410 | 800.00 |
| Bacteriocins-*FliC* | 0.088 | -1.138 | 1.403 | 800.00 |

| | | | | |
|---|---|---|---|---|
| *FliC*-T6SS | 0.074 | -1.188 | 1.410 | 800.00 |
| T6SS | 1.000 | 1.000 | 1.000 | 0.00 |
| Bacteriocins-T6SS | 0.000 | 0.000 | 0.000 | 0.00 |
| *FliC*-Bacteriocins | 0.088 | -1.138 | 1.403 | 800.00 |
| T6SS-Bacteriocins | 0.000 | 0.000 | 0.000 | 0.00 |
| Bacteriocins | 1.000 | 1.000 | 1.000 | 0.00 |

**Supplementary Table S9. Residual variance–covariance matrix (R structure) from the multivariate phylogenetic mixed model:** Posterior estimates of residual variances and covariances from the multivariate phylogenetic mixed model. Residual variances for threshold and count traits were fixed for model identifiability.

| **Trait** | **Posterior mean** | **Lower 95% CI** | **Upper 95% CI** | **Effective sample size** | **pMCMC** |
|---|---|---|---|---|---|
| *FliC* | 9.033 | -14.238 | 33.036 | 223.06 | 0.400 |
| T6SS | -3.639 | -4.728 | -2.351 | 800.00 | 0.001 |
| Bacteriocins | -0.524 | -0.781 | -0.312 | 800.00 | 0.001 |

**Supplementary Table S10. Fixed effects from the multivariate phylogenetic mixed model:** Posterior estimates of fixed-effect intercepts from the multivariate phylogenetic mixed model.

| **Parameter** | **Effective sample size** | **Autocorrelation (Lag 500)** |
|---|---|---|
| *FliC* phylogenetic variance | 2.78 | 0.974 |
| *FliC*–T6SS phylogenetic covariance | 4.39 | 0.756 |
| *FliC*–bacteriocin phylogenetic covariance | 586.69 | 0.059 |
| T6SS phylogenetic variance | 800.00 | -0.021 |
| T6SS–bacteriocin phylogenetic covariance | 800.00 | 0.024 |
| Bacteriocin phylogenetic variance | 800.00 | 0.001 |

**Supplementary Table S11. Convergence diagnostics for key phylogenetic variance–covariance parameters:** Effective sample size and autocorrelation diagnostics for key parameters from the phylogenetic random-effects structure. Low effective sample sizes and high autocorrelation indicate reduced chain mixing and increased uncertainty in parameter estimation.

| Analysis | N species/tips | 00 | 01 | 10 | 11 | Independent log ML | Dependent log ML | log BF |
|---|---|---|---|---|---|---|---|---|
| Pseudomonadota only | 1,711 | 364 | 91 | 678 | 578 | -1573.965 | -1565.625 | 16.68 |
| Excluding Pseudomonadota | 2,414 | 1627 | 65 | 670 | 52 | -987.098 | -995.014 | -15.83 |

**Supplementary Table S12. Clade-specific sensitivity analysis of the association between flagellar motility and T6SS presence.** BayesTraits analyses were performed within Pseudomonadota and after excluding Pseudomonadota from the 4,125-species phylogeny. The number of species in each combination of flagellar motility and T6SS presence is shown: 00, flagella absent/T6SS absent; 01, flagella absent/T6SS present; 10, flagella present/T6SS absent; and 11, flagella present/T6SS present. Log Bayes factors (log BF) were calculated as 2 × (dependent log marginal likelihood − independent log marginal likelihood).

| Transition | State of other trait | Mean | SD | Median | 95% CI |
|---|---|---|---|---|---|
| T6SS gain | Flagella absent | 119.0 | 34.4 | 114 | 65–197 |
| T6SS gain | Flagella present | 515.6 | 77.3 | 509 | 380–684.5 |
| T6SS loss | Flagella absent | 157.5 | 40.2 | 154 | 92.5–246 |
| T6SS loss | Flagella present | 617.7 | 78.1 | 612 | 480–787 |
| Flagella gain | T6SS absent | 25.4 | 8.1 | 25 | 11–43 |
| Flagella gain | T6SS present | 5.0 | 5.3 | 4 | 0–19 |
| Flagella loss | T6SS absent | 122.7 | 15.6 | 123 | 90–151 |
| Flagella loss | T6SS present | 62.8 | 14.2 | 62 | 39–94 |

**Supplementary Table S13. Stochastic character mapping of transitions between flagellar motility and T6SS presence within Pseudomonadota.** The mean, standard deviation (SD), median and 95% interval of the number of reconstructed gains and losses are shown across stochastic character maps for the Pseudomonadota-restricted phylogeny. T6SS gains and losses are reported according to whether flagella were present or absent, and flagellar gains and losses according to whether T6SS was present or absent.

| Transition | State of other trait | Mean | SD | Median | 95% CI |
|---|---|---|---|---|---|
| T6SS gain | Flagella absent | 16.2 | 6.9 | 16 | 5–30 |
| T6SS gain | Flagella present | 18.5 | 8.7 | 18 | 5–37 |
| T6SS loss | Flagella absent | 79.1 | 26.0 | 84 | 32–122 |
| T6SS loss | Flagella present | 97.1 | 16.5 | 99 | 64–126 |
| Flagella gain | T6SS absent | 74.0 | 7.2 | 74 | 60–89 |
| Flagella gain | T6SS present | 5.6 | 6.8 | 4 | 1–18.5 |
| Flagella loss | T6SS absent | 134.9 | 14.3 | 135 | 107–164 |
| Flagella loss | T6SS present | 23.0 | 9.6 | 23 | 7–43 |

**Supplementary Table S14. Stochastic character mapping of transitions between flagellar motility and T6SS presence after excluding Pseudomonadota.** The mean, standard deviation (SD), median and 95% interval of the number of reconstructed gains and losses are shown across stochastic character maps for the phylogeny excluding Pseudomonadota. T6SS gains and losses are reported according to whether flagella were present or absent, and flagellar gains and losses according to whether T6SS was present or absent.

| **Bacteriocin threshold** | **Bacteriocin-positive species (N)** | **Bacteriocin-positive species (%)** | **Dependent log ML** | **Independent log ML** | **log BF** |
|---|---|---|---|---|---|
| ≥1 HMM hit | 2552 | 61.9 | -3869.579 | -3879.298 | 19.44 |
| ≥2 HMM hits | 1187 | 28.8 | -3598.942 | -3617.366 | 36.85 |
| ≥3 HMM hits | 421 | 10.2 | -2560.988 | -2576.824 | 31.67 |
| ≥4 HMM hits | 124 | 3.0 | -1821.749 | -1815.295 | -12.91 |
| ≥5 HMM hits | 34 | 0.8 | -1450.375 | -1444.543 | -11.66 |

**Supplementary Table S15. Sensitivity of the association between flagellar motility and bacteriocin presence to the bacteriocin detection threshold.**
